# Temporal continuity shapes visual responses of macaque face patch neurons

**DOI:** 10.1101/2022.02.22.481478

**Authors:** Brian E. Russ, Kenji W. Koyano, Julian Day-Cooney, Neda Perwez, David A. Leopold

**Affiliations:** Section on Cognitive Neurophysiology and Imaging, National Institute of Mental Health; Bethesda, USA; Center for Biomedical Imaging and Neuromodulation, Nathan Kline Institute; Orangeburg, USA; Department of Neuroscience, Icahn School of Medicine at Mount Sinai; New York City, USA; Department of Psychiatry, New York University at Langone; New York City, USA; Neurophysiology Imaging Facility, National Institute of Mental Health, National Institute of Neurological Disorders and Stroke, National Eye Institute; Bethesda, USA

**Keywords:** Temporal Context, Inferior Temporal Cortex, Ventral Stream, onset responses, naturalistic viewing, movies

## Abstract

Macaque inferior temporal cortex neurons respond selectively to complex visual images, with recent work showing that they are also entrained reliably by the evolving content of natural movies. To what extent does visual continuity itself shape the responses of high-level visual neurons? We addressed this question by measuring how cells in face-selective regions of the macaque temporal cortex were affected by the manipulation of a movie’s temporal structure. Sampling the movie at 1s intervals, we measured neural responses to randomized, brief stimuli of different lengths, ranging from 800 ms dynamic movie snippets to 100 ms static frames. We found that the disruption of temporal continuity strongly altered neural response profiles, particularly in the early onset response period of the randomized stimulus. The results suggest that models of visual system function based on discrete and randomized visual presentations may not translate well to the brain’s natural modes of operation.

## Introduction

We experience the world as a continually evolving sequence of events, scenes, and interactions. Humans and other animals use their vision to track and integrate a wide range of information, coordinating the inherent dynamics of the natural world with their own active sampling of important scene elements (Melcher, 2001; Schroeder et al., 2010). This integration provides a predictive context for interpreting the flow of incoming visual signals. What role might high-level visual neurons, such as those observed throughout the macaque inferior temporal cortex, play in such integration? For example, would cortical neurons apparently devoted to the processing of faces be influenced if the normal continuity of visual experience were artificially broken up into discrete, randomized presentations?

It is difficult to make predictions on this point, since nearly all that is known about visual responses in the brain derives from highly reductionistic modes of stimulus presentation. Traditional studies have used the abrupt presentation of isolated visual images to demonstrate object selectivity among neurons in the macaque inferior temporal cortex (Desimone et al., 1984; Gross et al., 1972), often for ethologically relevant categories such as faces and bodies (Jellema and Perrett, 2003; Perrett et al., 1992; Tsao et al., 2006). By comparison, a few single-unit studies have investigated how such neurons respond under more naturalistic conditions, for example when a subject is allowed to freely gaze within an evolving visual scene (McMahon et al., 2015; Mosher et al., 2014; Sheinberg and Logothetis, 2001). Thus, much more is known about how the brain responds to flashed presentations of isolated stimuli than how it responds under the continuity of everyday vision.

The investigation of visual temporal sequences has been pursued in some areas, such as the brain’s encoding of bodily actions (Jellema and Perrett, 2003; Keysers and Perrett, 2004; Nelissen et al., 2005; Singer and Sheinberg, 2010; Vangeneugden et al., 2009). These studies have linked the comprehension of bodily actions to the integration of form, movement, and location (Jellema et al., 2004), and in some cases to the generation of one’s own bodily actions (Keysers and Perrett, 2004). Reductionistic investigations of sequence processing in the inferotemporal cortex has in some cases identified a marked temporal dependency, which has often been linked to Hebbian or other forms of statistical learning through repeated sequence presentation (Meyer and Olson, 2011; Meyer et al., 2014a; Miyashita, 1988; Perrett et al., 2009; Ramachandran et al., 2017; Schwiedrzik and Freiwald, 2017). These studies provide evidence that object-selective neurons in the inferior temporal cortex exhibit significant temporal dependency in their responses.

A somewhat different facet of temporal integration involves the active sampling of a visual scene provided by the frequent redirection of gaze (Henderson, 2003; Leszczynski and Schroeder, 2019; Schroeder et al., 2010). For humans and other primates, whose high-resolution vision is sharply concentrated in the fovea, the sequential gathering of important information across fixations is a central feature of visual cognition (Mitchell and Leopold, 2015). In normal vision, the temporal discontinuities introduced by this active sensing of the visual scene through gaze redirection is superimposed upon the temporal dynamics of actors and events in the external world. The stable perception of the world amid the constantly shifting retinal stimulation is one of the enduring puzzles of visual neuroscience (Efron, 1967; Melcher and Colby, 2008; Wurtz et al., 2011).

Recently, the use of naturalistic stimulus paradigms has gained popularity and has begun to complement more conventional presentation methods (for a recent review, see (Leopold and Park, 2020)). The human fMRI community has, in particular, taken up the use of movies as a new paradigm for tapping into various aspects of brain function (Bartels and Zeki, 2004; Çukur et al., 2013; Franchak et al., 2016; Hasson and Honey, 2012; Hasson et al., 2004, 2008, 2010; Huth et al., 2012; Vanderwal et al., 2015; Vodrahalli et al., 2017; Wang et al., 2012), with some monkey investigators following suit (Mantini et al., 2012, 2013; Park et al., 2017; Russ and Leopold, 2015; Russ et al., 2016; Shepherd et al., 2010; Sliwa and Freiwald, 2017). One single-unit study investigated responses in macaque anterior fundus face patch to repeated 5-minute presentations of naturalistic movies full of dynamic social interaction (McMahon et al., 2015). Neurons in this area were strongly entrained by the movie timeline, with a high level of response repeatability across presentations of the same movie despite the free movement of the eyes. However, the role of temporal continuity in shaping neural responses to complex movie content is at present poorly understood. While not yet studied in detail, this question may be important, since our understanding of high-level vision is at present shaped strongly by the brain’s response to the presentation of discrete and isolated stimuli.

The current study addresses this question using a paradigm in which we systematically manipulated the temporal continuity of visual content stemming from a naturalistic movie. We recorded from single neurons in the anterior fundus (AF) and the anterior medial (AM) face patches and compared spiking responses to continuous versus discrete presentations of the same movie content. We found that the artificial disruption of temporal continuity significantly affected neural responses to the movie content in both areas. The strongest impact was on the response to the initial onset of the discrete presentation, for which neural responses were nearly uncorrelated to those observed in response to the same content of the intact movie.

## Results

Five macaque monkeys (*Macaca mulatta*) viewed intact movies and randomized movie clips (“snippets”) as visual responses were recorded from single neurons in the anterior fundus (AF) and anterior medial (AM) face patches. These regions were defined functionally using fMRI (Tsao et al., 2008) and targeted using methods described previously (McMahon et al., 2014b, 2014a) (**Figure 1a**, see **Methods**). We recorded from 108 neurons in the AF face patch (Subject T: 18; Subject R: 16; Subject S: 74) and 72 neurons in the AM face patch (Subject D: 49; Subject M: 23). For the main analyses, we combined neurons across regions into a single population after determining that the results were similar in both regions. Corresponding analyses performed separately in the two face patches are presented in supplemental material. The animals viewed multiple repetitions of an intact 5-minute movie. They also viewed randomized short snippets and static images extracted from the movie at each of 300 time points spaced by one second (see **Methods**). Most of the analyses described below involve comparing the responses to 300 equivalent visual segments from the original, intact movie with either 800 ms movie snippets or 100 ms static images. Corresponding responses to two additional conditions (250 and 100 ms movie snippets) are also discussed.

**Figure 1.**
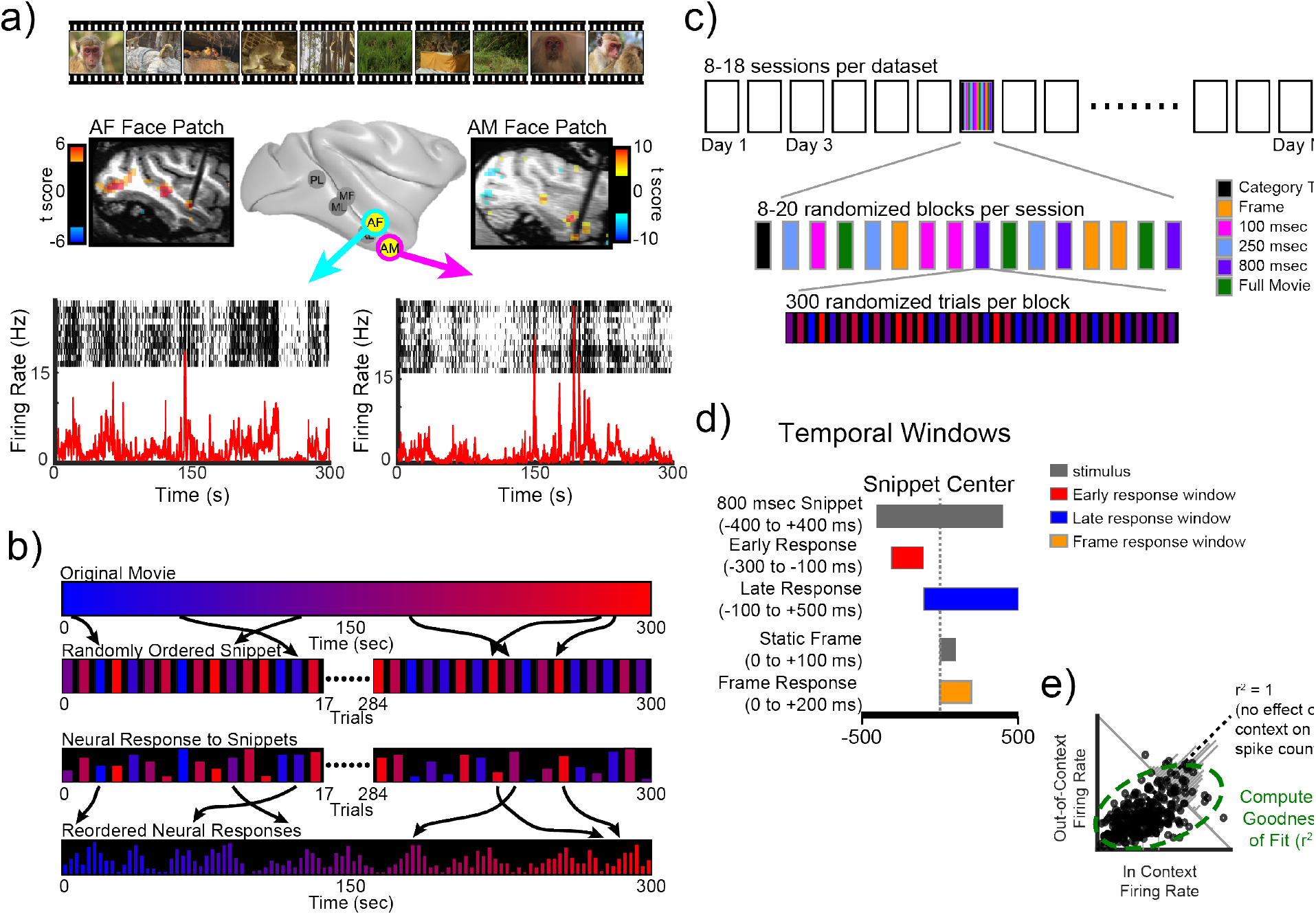
Experimental Design. A) Top row shows representative frames from the intact movie that was presented during the free viewing paradigm. Middle row depicts the results of the fMRI face-patch localizer task used to target the AF (left) and AM (right) face patches. Functional localizer is overlaid on a T1w image taken after the electrodes where implanted (see Methods). The bottom row depicts the spiking response from one AF (left) and one AM (right) neuron to the viewing of multiple trials of the intact movie. The black dots represent the time of a spike with 1 ms bins, and each row is a trial. The red line is the mean spike density function for all trials. B) Schematic of the creation of the individual snippets from the original movie, and the reconstructed response to those snippets. C) Schematic of the pseudorandomized task design that was used to collect the data within and across sessions. D) Schematic for the selection of response periods based on the individual snippet lengths. E) Schematic of the regression analysis between the snippet responses and the original movie. The green circle represents a hypothetical goodness of fit measure.

The recordings hinged on the longitudinal recording capabilities of the implanted microwire bundles since the data for individual neurons was amassed over 8-18 daily sessions of data collection (McMahon et al., 2014a, 2015)(see **Fig S1**). Each of these sessions began with several minutes of conventional testing with flashed images from different stimulus categories, which was used both to establish each neuron’s basic selectivity and as a tool for “fingerprinting” to identify the same neurons across subsequent recording sessions. This testing was followed by all other conditions presented in randomized blocks (**Figure 1c**). The intact movie was viewed between 8 and 24 times (median for viewings per neuron: monkey T = 8, monkey R = 10, monkey S = 15, monkey M = 24, monkey D = 18). The randomized movie snippets were viewed between 7 and 37 times each (median viewings per neuron: 800 ms: T = 7, R = 10, S = 13, M = 13, D = 12; 250 ms: T = 7, R = 11, S = 16, M = 14, D = 17; 100 ms: T = 7, R = 9, S = 17, M = 13, D = 20; static image: T = 13, R = 17, S = 37, M = 33, D = 28). The following sections describe the consequences of random ordering and flashed presentation on the responses of neurons to the visual content of a naturalistic movie.

### Temporal context affects responses to movie content

Neural responses to movie content were strongly affected by the continuity of the video. **Figure 2** shows the responses of an example neuron (neuron AF094) to the same content in continuous versus randomized presentation. This neuron was typical in that it was face-modulated and entrained reliably by the movie. **Figure 2a** shows the raw raster responses during representative 800 ms snippets and corresponding static frame presentations. From these example rasters, the repeatability across presentations and the effects of randomization are both evident. **Figure 2b** shows the reconstructed time courses to the 800 ms snippets early onset responses (red, 100-300 ms following snippet onset) and the later responses (blue, 300-900 ms following snippet onset) in comparison to the equivalent time periods from the continuous movie. The responses to the randomized presentation are reordered to match the native timeline of the movie (see **Figure 1b**, **Methods**). Both reconstructed time courses revealed notable deviations in spiking during the corresponding analysis windows of the continuous presentation (gray time courses). **Figure 2c** applies a similar analysis to spiking responses to the static single frames drawn from the movie, which showed little correspondence to the spiking activity in the continuous condition. Scatterplots showing the response correspondence for the 300 sample time points are shown to the right of each reconstruction. For this example neuron, the initial onset responses to both the snippet (red) and static frame (yellow) were uncorrelated with the responses to the equivalent visual content presented in the continuous condition (snippet: r = 0.0665, p = 0.2506; static frame: r = −0.031, p = 0.5933). The responses to content later in each snippet (blue) were affected by randomization, but retained their correlation with the continuous responses (r = 0.55, p = 0.01e-23). Additional single unit examples from both areas are shown in **Figure S2**. Across the population, two principal factors governed the influence of randomized snippet presentation on neural firing. The first factor was the abrupt onset of the stimulus and the second was the temporal reordering of the movie content. We investigated the expression of these two factors across the neural population.

**Figure 2.**
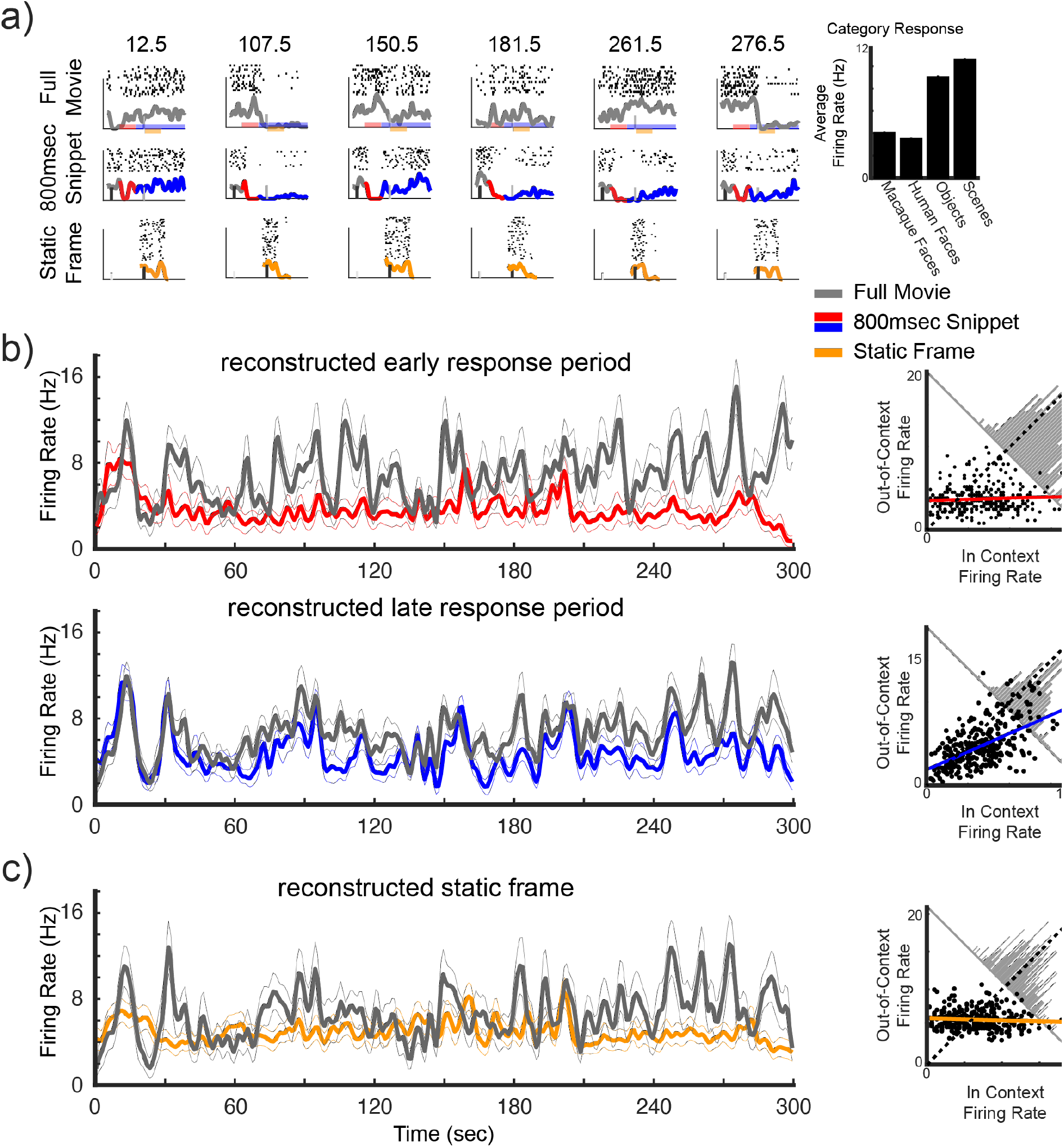
Representative response of an AF neuron. Six representative time points are featured (A), which show the diversity of response across time points and contexts. We compared stimulus-driven response during the intact movie (top row), 800 ms snippets (middle row) and 100 ms static centerframe (bottom row). The rows of spiking data show the raster plots and computed average spike density function during the three conditions for the featured time points. The mean firing rate (Hz) to the static categorical images is presented next to the snippet responses, highlighting the current neurons face responsiveness. B) shows average intact movie response (gray) as well as the reordered spike count responses to the transition response (red top), the static frame (orange middle), and delayed response (blue bottom) using matching analysis windows relative to the center frame (see Figure 1b,d).

### Abrupt onset strongly alters visual responses

Across both face patches, the early onset response selectivity to either a snippet or static frame was often statistically uncorrelated with that measured during the native movie presentation. To further quantify this relationship, we applied linear regression to the 300 samples collected in the corresponding conditions (see scatterplots in **Figure 2**), providing a measure of goodness-of-fit (r^2^) for each comparison. **Figure 3a** shows that, across the population, the initial onset response to the 800 ms snippets had a poor correspondence to the visual responses elicited during the continuous condition (r^2^_mean_ = 0.11 +/- 0.12). This was a sharp drop from the split-halves control condition (black), in which the same number of trials was used to compute within-condition r^2^ values (paired t-test(103)= 25.99, p = 0.5e^-48^). The low correspondence to the native responses was even more evident when static frames were used for the comparison (**Figure 3b,** r^2^_mean_ =0.06 +/- 0.08). Compared to the within-condition correspondence, this also exhibited a large and significant decrease (**Figure 3b**, paired t-test(103) = 22.18, p = 0.5e^-42^). These findings demonstrate that the abrupt appearance of a static image or dynamic clip on a blank background elicits initial onset responses that differ markedly from the responses to the same content presented within the context of the movie (see **Figure S3** for analyses for separate patches). In addition to the 800 ms movie snippet featured above, analysis of the onset period for the independent 250 ms and 100 ms snippet presentations yielded similar results (**Figure S4**).

**Figure 3.**
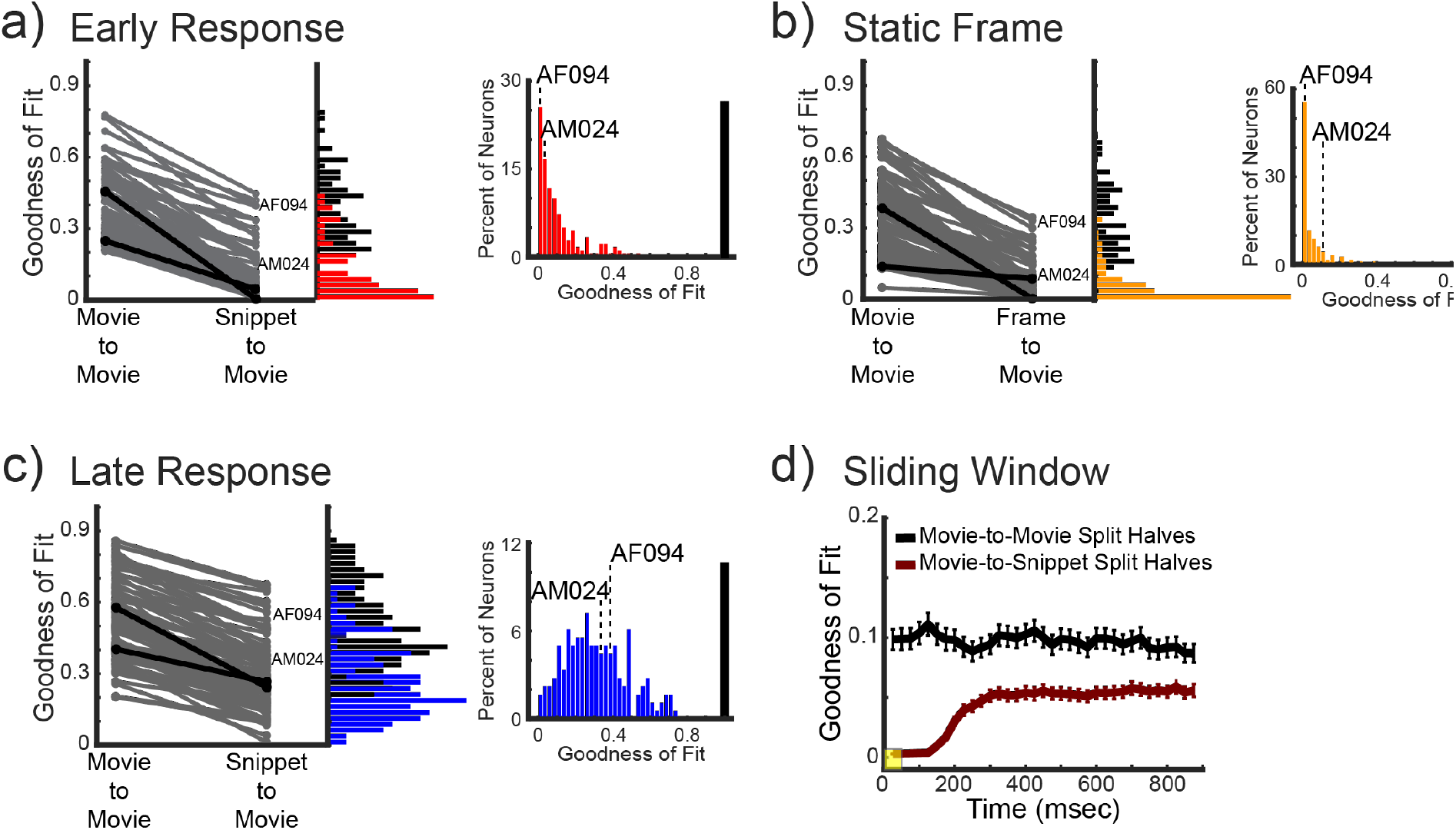
Population Response changes across context and time. A) Paired comparison within context and between context goodness-of-fits (r^2^) of neurons in the AF and AM face-patches during the onset period. Neurons in the pair comparison were filtered by an r^2^ or 0.2 in the within-context condition. The black lines highlight the example neurons presented in **Figures 2** and **S2b**. The vertical histograms represent the density of r^2^-values for the split halves within-context (black) and between context (red) for the full population. The lower histogram shows the r^2^ for all trials of the between context regression for the full population. B) Shows the same set of comparisons for the response to the presentation of the static frames. (C) depicts the same data and analyses for the late period of the 800 ms snippets. (D) show the evolution of the goodness-of-fit for the within-context (black) and between context (red) regressions using a sliding window of 50 ms (yellow box) with 25 ms jumps. Error bars represent the standard-error of the mean of the population within each bin.

### Temporal continuity shapes visual responses

During the later period of the 800 ms snippets, beyond the neurons’ transient response to the initial stimulus onset, it was possible to assess the effect of temporal continuity itself without the contribution of the onset response period. As before, responses were compared during periods of equivalent stimulus content in the native movie and in the snippets. During the snippet, this pertained to a 600 ms time window beginning 300 ms after the onset.

These late period responses of face patch neurons varied in their level of disruption by temporal randomization, with some neurons moderately affected by temporal reordering (e.g. **Figure 2**, **Figure S2a**) and others showing little if any influence (e.g. **Figure S2b**, **Figure S2c**). Across the combined AF/AM population, the influence of temporal continuity was evident as a broad distribution of r^2^ measures (**Figure 3c**, r^2^_mean_ =0.31 +/- 0.17), which contrasted with the narrow r^2^ distributions of the early responses (**Figure 3a**). Separate analyses for the AF and AM face patch neurons are provided in **Figure S3**. As above, the r^2^ values were contrasted with those obtained from a within-session split-halves control condition. The r^2^ values for the late responses were significantly higher than for the early responses (2×2 (Context x Time Frame) ANOVA: main effect of Context F(1,716)=233.04; p=0.9e^-47^; Main effect of Time Frame F(1,716)=162.24 p=0.1e^-34^; but no interaction F(1,716)=0.054; p=0.82). Nonetheless, the temporal reordering had a highly significant effect well beyond the initial response onset (paired t-test(148)= 30.78, p = 0.3e^-66^). Thus, temporal continuity itself contributes in an important way to the responses of face patch neurons during video viewing.

We further examined the temporal dynamics of the response decorrelation that accompanied the initial onset of the randomized stimuli. The traces in **Figure 3d** track the r^2^ values using a running window (25 ms jumps) for both in context (movie-to-movie) and out of context (movie-to-snippet) comparisons. Note that the absolute r^2^ values are lower in magnitude than before owing to the narrow (50 ms) bin width of the analysis (see **Methods**). The results revealed that the profound disruption of visual responses began to subside at approximately 200 ms following the presentation of the snippet. Together, these results demonstrate two clear response phases following the onset of the movie snippets: namely, an early onset period in which response selectivity is severely disrupted, and a later period in which the temporal shuffling affects the responses of some neurons strongly and others not at all.

### Relationship to face selectivity

While the face selectivity of the neurons did not figure prominently into the design of the present study, we did confirm that many of the neurons we tested in and around the AF and AM face patches exhibited a preference for faces over other images (**Figures 4a; Figure S5**), in accordance with many previous studies (Aparicio et al., 2016; Bell et al., 2011; Freiwald et al., 2009; Popivanov et al., 2014; Tsao et al., 2006). Specifically, we found that 88 of the 180 recorded neurons, or 49% of our population, had a *d’* greater than 0.65 which is a criterion similar to that used by previous studies (Aparicio et al., 2016; Popivanov et al., 2014). To test whether a cell’s category selectivity had bearing on its observed dependency on temporal context during naturalistic stimulus presentation, we compared each neuron’s d’ score for face selectivity to the magnitude of its between context goodness-of-fit values. The results indicated a significant, though weak, relationship between the face selectivity d’ value and context r^2^ value in the early response period (Pearson’s correlation = 0.1716, p = 0.0213). Thus, the more highly face selective neurons showed a higher retention of visual selectivity during the initial onset response. Importantly, this relationship was not present during the later response period (Pearson’s correlation = 0.0738, p = 0.3248), where the face selectivity of a neuron was unrelated to its resilience to temporal randomization.

**Figure 4.**
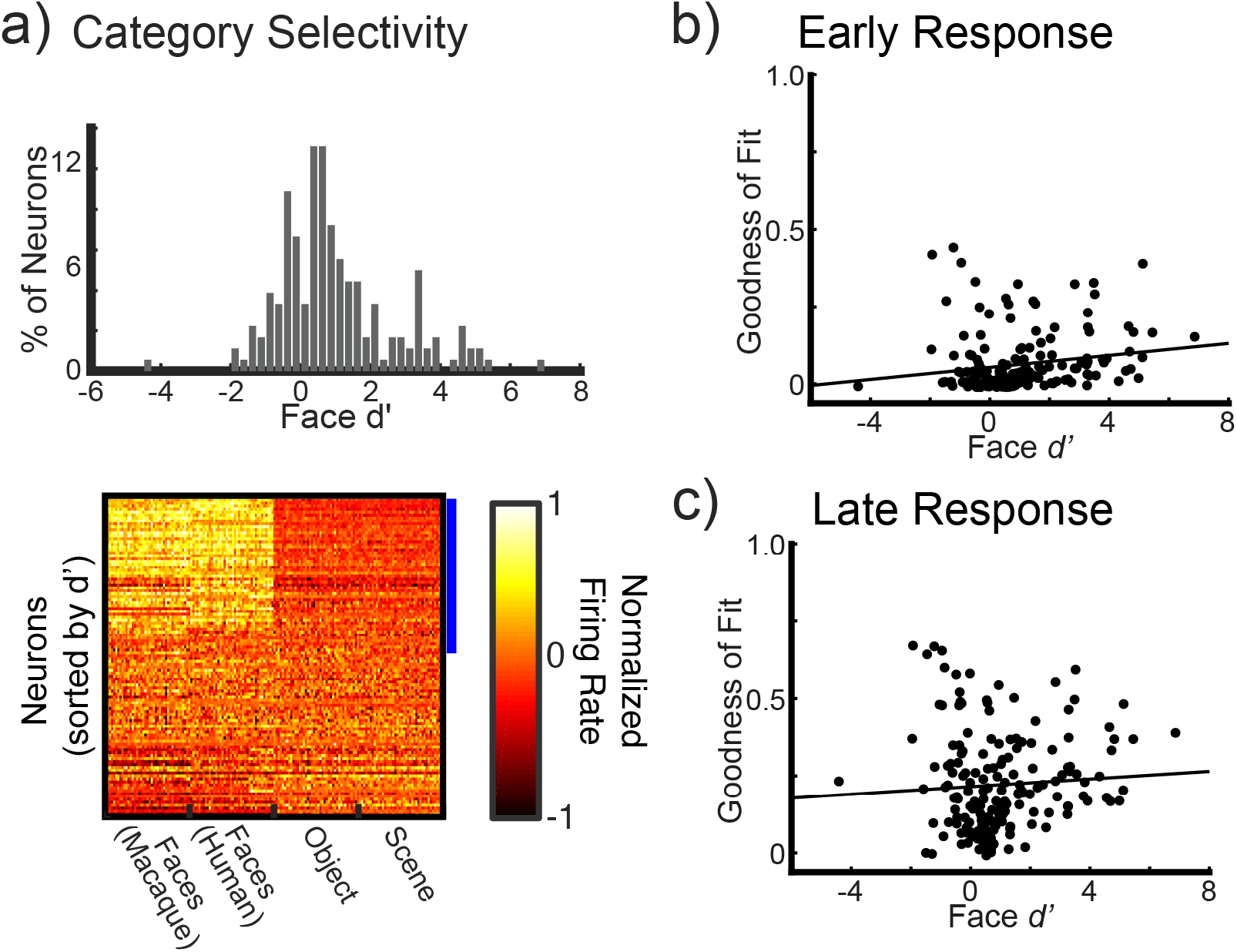
The interaction between face selectivity and context responses. (A) Selectivity of the combined face patch neurons to faces over scenes and objects. The upper histograms depicting the density of d’ values for all neurons. Values of d’ greater than zero represent a preference for larger responses to faces. The lower heat maps show normalized firing rate, to the maximum response of a given cell, for all neurons within the populations sorted by their d’ score. The blue bar on the side of the heat map shows neurons that had significant d’ values, and thus would be categorized as face selective. (B) and (C) show the relationship between face selectivity and our goodness-of-fit (r^2^) measure for the early and late response periods, respectively. The black line shows the line of best fit between the two response measures.

## Discussion

In the current study, we found that the responses of neurons in anterior face-selective regions of the macaque inferior temporal cortex differ depending on temporal context. Neurons responded differently to the same movie content when it was presented continuously versus when it was divided into brief snippets and shown out of context, in randomized presentation. The disruption in visual responses was particularly prominent in the period shortly following the onset of the snippet. In the following sections, we discuss how visual operations may differ between the continuous, free viewing of a movie and the brief, randomized presentation of its content. We also discuss the role of perceptual history in the establishment of spatial and temporal context during normal vision, as well as the bearing of these results on the activity of face-selective neurons.

### Continuous vs. discrete stimulus viewing

The change in responses observed for the same movie content during continuous versus randomized presentation may, in part, reflect a difference in the way in which the brain is prepared to process stimuli in different visual contexts. Most visual neurophysiology experiments explicitly use discrete presentation to show stimuli that are isolated in space and time. This has led to a view of the object processing pathway in which neurons in the inferior temporal cortex analyze visual form in a way that depends little on temporal or behavioral context. Some work, however, has suggested that where an animal is looking (DiCarlo and Maunsell, 2000, 2003; Sheinberg and Logothetis, 2001) or what an animal has recently seen (Singer and Sheinberg, 2010) can strongly affect the responses of IT neurons. Such neurons have been shown to modify their responses based on contextual information, usually in the form of a cue or memory component (Anderson et al., 2008; Kourtzi and DiCarlo, 2006; Kuravi and Vogels, 2017; Meyer et al., 2014b; Miller et al., 1991; Rolls et al., 1989; Sheinberg and Logothetis, 2001; Vinken et al., 2018; Woloszyn and Sheinberg, 2012). Neurons in the superior temporal sulcus are associated with motion stimuli (Furl et al., 2012; Nelissen et al., 2005, 2006, 2011; Polosecki et al., 2013; Vangeneugden et al., 2009), in particular biological motion, for which temporal continuity may be particularly important. Increasingly, researchers are using naturalistic stimuli as a complementary approach to conventional stimulus presentation (for a review, see (Leopold and Park, 2020)).

The majority of studies investigating temporal contributions of IT neurons have not attempted to quantify the time scales over which neurons integrate visual information. In the current study, we attempted to address this by systematically varying the amount of temporal contextual information that was presented, in the form of brief movie snippets lasting 800 ms, 250 ms, 100 ms snippets, and static frames presented for 100 ms, in each case extracted from a longer movie. We found that the majority of neurons within both the AF and AM face patches appear to integrate temporal information over extended time periods, since even the late period responses of the shuffled snippets always differed from the responses to the native movie, as verified by analysis controlling for equivalent numbers of trials (**Figure 3**).

Within higher-order ventral stream regions there is little research specifically addressing temporal integration scales at the neuronal level, there is evidence of similar integration at the regional level using fMRI. Hasson et al (Hasson et al., 2008) used a naturalistic viewing paradigm somewhat similar to the one we implemented to investigate the length of temporal receptive fields throughout the brain. In a set of fMRI studies, they found that temporal receptive fields, the length of coherent temporal context necessary to engage a region, increased systematically as information moved from early visual cortex through the visual system into the prefrontal cortex. In relation to the current study, they found that regions in the temporal lobe of their human subjects began to engage similarly when the subjects viewed 4-8 seconds of coherent temporal information. While these data were obtained using fMRI, which has a much slower timescale than single-unit activity, and a slightly different experimental paradigm than the current study, they support our results in suggesting that neurons within the temporal lobe may incorporated greater than one second worth of contextual information in determining their response to a given stimulus. In future research, one will need to expand the range of temporal contexts used to more fully map the temporal space within these regions.

### Onset disrupts visual responses

The largest difference between responses was observed during the initial phase of the response to randomized stimuli, following the abrupt onset of the stimulus. Specifically, during the first 200-300 ms of randomized snippets and static frames, the responses had virtually no correspondence to the full movie responses. While this result may be expected in some ways, as onset responses have been shown to cause nonspecific transients throughout the visual system (Hung et al., 2005; Müller et al., 2001; Oram and Perrett, 1992; Schiller and Malpeli, 1977), it is critical to note that the majority of previous experiments that probe feature and category response live in these early onset responses. Additionally, while the early and static frame responses exhibited low correspondence with the original context, it was not simply due to the strong onset excitability during this period. For some neurons and snippets, the out-of-context early period was relatively silent, whereas in the full movie a higher firing rate was observed (e.g., response at 150.5 s in Fig.2a). Nor is it due to a large delay in response onset in these neurons. The average response latency for both populations (AF = 115 ms; AM = 120 ms), as calculated relative to the static frames onset, was well before we saw the plateau in the correspondence between the out of context response and the native movie, and more important was within the time period analyzed for the early response window. These variable responses to the snippets during the early phase suggest that differences in response are not simply due to non-specific onset response, but instead potentially due to a lack of full contextual information for the stimulus, which has been neglected in most conventional visual experiments.

### Rethinking the role of feature-selective neurons in natural vision

We found that face selectivity had little relation to a neuron’s level of disruption by temporal randomization, though there was a significant positive correlation between a neuron’s degree of face selectivity and its preservation of response selectivity during the early onset phase of randomized presentations. The strong influence of continuous temporal structure upon the responses of face selective neurons underscores the concern that conventional modes of testing only capture a narrow range of visual function in object processing region of the cortex (Leopold and Park, 2020; Park et al., 2017, 2021). It is interesting to consider that, during periods of visual free viewing, inferior temporal cortical neurons, including those face patch neurons reported here, enter into a mode of operation marked by sequential target selection, continuity of visual content and attentional processes, and anticipation of events. In this exploratory mode, and in contrast to discrete modes of image presentation, basic visual responses may be modified by neuromodulatory inputs, along with a range of brain areas involved in memory, spatial perception, motor planning, and executive function. As experimental work continues to explore the frontiers of natural visual experience, our portrait of visual brain organization may need to evolve and expand. A central feature of natural experience that is explicitly removed from most experimental approaches is that of temporal continuity. The present study demonstrates that, for populations of well-studied feature selective neurons in the macaque high-level visual cortex, temporal continuity critically determines visual response profiles.

## Methods

### Subjects

Five rhesus macaques (*macaca mulatta*) participated in the current experiments, including 2 females (ages 4-13 years at the time of the study) and 3 males (ages 4-10 years). All procedures were approved by the Animal Care and Use Committee of the National Institutes of Mental Health, and followed the guidelines of the United States National Institutes of Health.

In an initial surgery, each animal was implanted with a fiberglass head-post, for immobilizing the head during recording sessions. After behavioral training, described below, microwire bundle electrodes (Microprobes) and adjustable microdrives were implanted into the face patches during a second surgical procedure (see **Figure 1a**). The electrodes, microdrives, and surgical procedures are described in a previous publication (McMahon et al., 2014b, 2014a). In three subjects (two female, one male) the electrodes were directed to the anterior fundus face patch (AF): Subject T received bilateral 32 channel electrodes bundles; Subject R received a 64 channel bundle in the right hemisphere; and Subject S received a 64 channel bundle in the left hemisphere. In two other subjects (both male), electrodes were directed to the anterior medial (AM) face patches: Subject M received one 64 channel microwire bundle in the left AM; Subject D was implanted bilaterally with 128 channel microwire bundles, with the left bundle located within AM and the right bundle just adjacent to AM.

Following electrode implantation, the subjects participated in a series of behavioral tasks. During this period, the subjects’ access to liquid was controlled, such that they were motivated to receive liquid reward in exchange for participating in the experimental tasks. In the case that they did not receive sufficient liquid reward in a given session, they were supplemented with liquids in their home cage. Hydration and weights were monitored throughout the experimental period to ensure the animals’ health and well-being.

### Visual Stimuli and Behavioral Monitoring

Subjects viewed a range of intact movies, randomized snippets and static frames extracted from the movies, and flashed categorical images. Image presentation and gaze fixation monitoring were under computer control using a combination of custom written QNX (Sheinberg and Logothetis, 2001) and Psychtoolbox (Brainard, 1997) code. Neurophysiological signals were recorded using TDT hardware and software. The subjects’ horizontal and vertical eye position was continuously monitored using a 60 Hz Eyelink (SR Research) system, which converted position to analog voltages that were passed to the neurophysiological and behavioral recording software. At the start of each session, the subject’s eye position was centered and calibrated using a nine-position grid with 6° visual angle separation. This calibration procedure was rerun repeatedly throughout each session to ensure good calibration during all experimental trials. Blocks of intact movie presentation, randomized snippet presentation, and categorical image presentation were randomly interleaved (**Figure 1c**) and took the following form.

#### Intact Movie Presentation

The monkeys repeatedly viewed multiple repetitions of a 5 min movie containing a wide range of interactions, mostly between other monkeys (McMahon et al., 2015; Russ and Leopold, 2015; Russ et al., 2016). The movies were presented on an LCD screen with a horizontal dimension of 15 dva. While there was no fixation, subjects were required to maintaining their gaze within the movie frame, for which they received a drop of juice reward each 2 − 2.5 s.

#### Randomized Snippet Presentation

The content from the movie above was broken down into short movie snippets and static frames, which the monkey viewed in randomized order. The original 5m movie was first divided into three hundred 1 s segments, which were subsequently used to generate snippets of different lengths. The snippet durations of 800, 250, and 100 ms spanned the center frame of each 1 s movie segment. For example, the 800 ms snippet from the 151^st^ second would present the frames of the movie from 150.10 to 150.90 s, whereas the 100 ms snippet from the same segment would display the frames from 150.45 to 150.55 s. The static frame condition presented the center frame for 100 ms. For each snippet length and the static frame condition, all 300 stimuli were presented repeatedly and in randomized order (**Figure 1b**). Other aspects of the stimulus were the same as in the intact movie condition, other than the 200 ms blank period now inserted between each presentation. As above, the subjects were permitted view the entire snippet content within the brief presentation period and were rewarded with a drop of juice and a 1.0 s break after completing approximately 2 s of trials, depending slightly on the snippet lengths. For an inappropriate fixation break, the animal would forfeit its juice for that period and incur an additional 750 ms timeout followed by repetition of the previous stimulus. For each block of the snippet task, the stimulus duration was held constant until all 300 stimuli were presented.

#### Categorical Image Presentation

In these blocks, the monkeys serially viewed brief image presentations drawing from four categories, with 40 images from each category. The image categories were monkey faces, human faces, objects, natural scenes. The stimuli were randomized and presented at 12 dva for 100 ms with a 200 ms inter-stimulus interval. As above, the subjects were permitted to direct their gaze anywhere within the image. If gaze was maintained in the window for seven successive images, the subject received a liquid reward and a 1000 ms inter-trial break. An inappropriate fixation break led to forfeit of reward and an additional 750 timeout. This condition served the dual role of evaluating category selectivity and establishing a “fingerprint” for tracking the identity of single neurons across sessions (McMahon et al., 2014b).

### Longitudinal Neural Recording

Given the large number of conditions and presentations required in the current study, our recording paradigm hinged on the capacity to accurately record the same neurons across multiple sessions. This longitudinal tracking of individual cells required particular care in spike sorting and rules for determining the continuous isolation of neurons across sessions.

#### Spike Sorting

Following each session of recording, channels were cleaned and then submitted to an automatic spike sorting algorithm. The cleaning procedure, run separately on each bank of 16 channels sharing a connector, involved regressing out the first principal component across channels. This procedure was important in our experience to combat certain types of movement and chewing-related artifacts that were shared across channels. Following this cleaning procedure, the automatic sorting program wave_clus (Chaure et al., 2018; Quiroga et al., 2004) was applied to identify and sort action potentials on individual channels. Candidate spike waveforms were first marked as any voltages exceeding four standard deviations of the noise. These candidate waveforms were then sorted automatically using PCA space, rendering a sequence of timestamps for between 0 and 4 usable waveforms from each microwire channel.

#### Tracking Neurons Across Days

Verifying the isolation of a given neuron across days required a combined consideration of spike waveform, as determined by the automated spike sorting algorithm (Chaure et al., 2018; Quiroga et al., 2004), and an independent comparison of the response fingerprint, or selectivity to the flashed images (see **Figure S1** for example of isolation and fingerprinting). A cell was determined to be the same as that isolated on a previous day if it (1) was recorded from the same channel, (2) had the same fingerprint signature of response selectivity, and (3) had a similar waveform. Of these three criteria, the third was the only one that was subject to flexibility, since it is well known that small positional changes in the relationship between an electrode and a neuron can change the action potential shape. However, any neuron recorded from a different electrode or having a different selectivity fingerprint across days were considered to be a different neuron. Applying these criteria, we excluded from analysis neurons that were held for fewer than 75% of the sessions, and therefore had insufficient numbers of trials for our statistical analyses. All visually responsive neurons, defined as those giving statistically significant responses in any one of the conditions, were investigated for the effects of temporal continuity, with a separate analysis investigating the specific factor of face selectivity.

### Data Analysis

The main data analysis focused on comparing the neural responses to the intact movie with those elicited by the same visual content presented during the randomized 800 ms snippets. Based on pilot analysis, we divided this analysis into two distinct time windows (**Figure 1d**). The first analysis window analyzed spiking from 100-300 ms following the onset of the snippet, targeting the contribution of the initial response. The second analysis window then focused on the delayed, or extended, response between 300-900 ms.

For each time window, we compared the mean spike count over multiple presentations of the 300 snippets to the corresponding mean spike counts elicited by the corresponding frames in the intact movie. In the main analysis, we compared the two conditions using a linear regression, from which we derived the goodness of fit (r^2^) to assess the effects of context (see **Figure 1e**). We also reordered the randomized snippet responses into their original sequence in order to visualize any temporal disruption (see **Figure 1b**).

For the main analysis, the r^2^ parameter provided insight into the disruption in visual selectivity introduced by randomized snippet presentation for the two time windows. For example, r^2^ values of 1.0 would indicate that a neuron exhibits identical responses across the two presentation conditions, thus the temporal context has no bearing on neural selectivity, whereas an r^2^ value of 0.0 would correspond to a complete disruption of the stimulus selectivity. For intermediate values, the r^2^ indicates the tightness of the stimulus selectivity relationship across conditions, or response variance explained.

In addition to the 800 ms snippets, we also analyzed neural responses to equivalent snippets of 250 and 100 ms, as well as static images of the center frame presented for 100 ms. In those cases, the presentation times were too short to allow for a separate consideration before and after the initial stimulus onset, thus a standard response window of 50 to 250 ms was used for evaluation of the onset response.

## Acknowledgements

This work was supported by the Intramural Research Program of the National Institute of Mental Health (ZIAMH002838, ZIAMH002898) to D.A.L.

## Supplemental Figures

**Figure S1.**
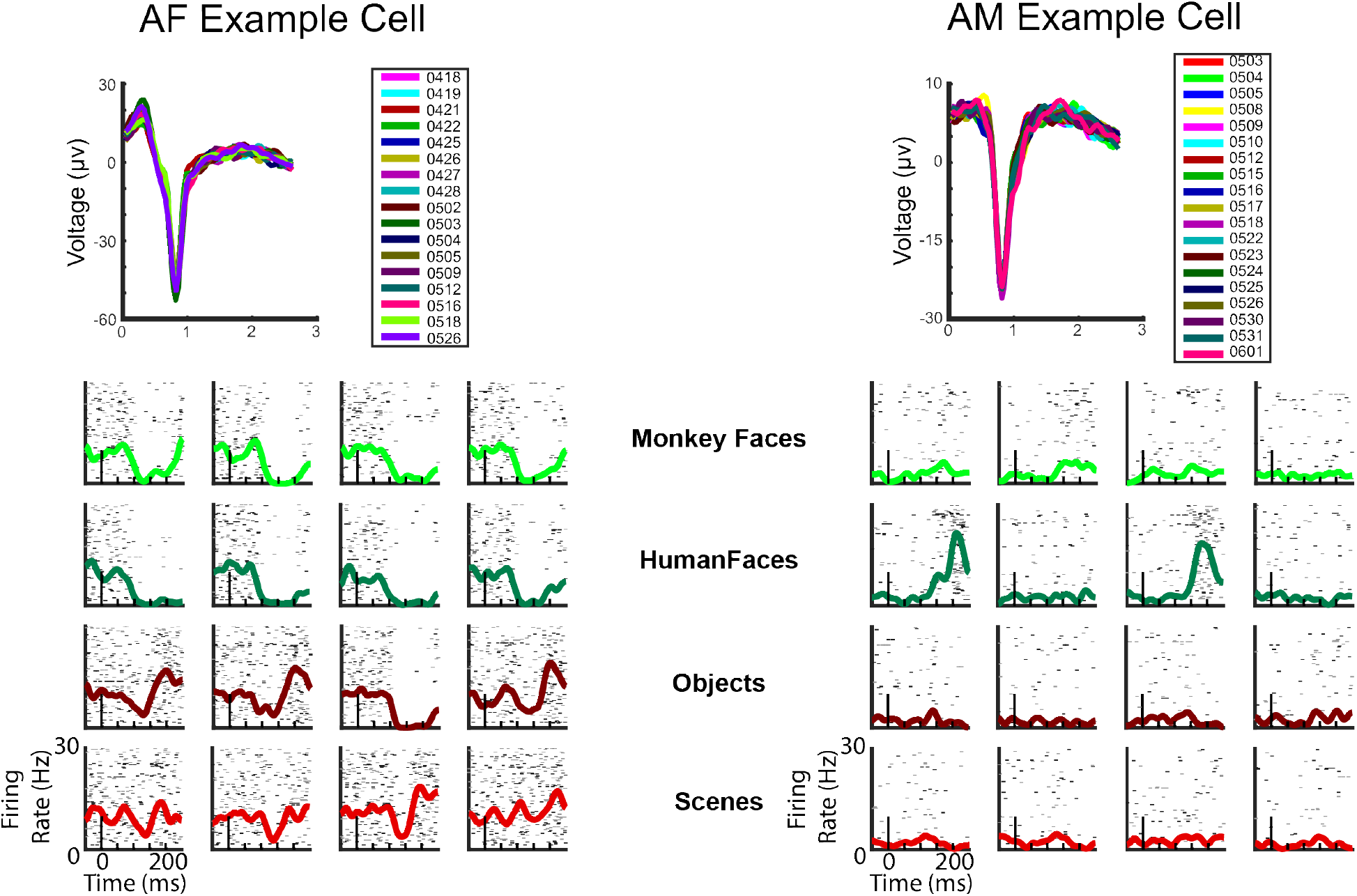
Example neurons from AF and AM after multisession sorting. (A) Shows the average shape of the sorted spikes for an example neuron from both AF and AM. Each color is represents the average spike waveform from a given session. (B) Shows the same neurons’ category response across multiple sessions for 4 example stimuli from the four categorical classes (Monkey Faces, Humans Faces, Objects, and Scenes). Each row represents a trial, and the color of trial alternate between grey and black based on session. The colored line shows the average spike density function for all trials collapsed across all sessions.

**Figure S2.**
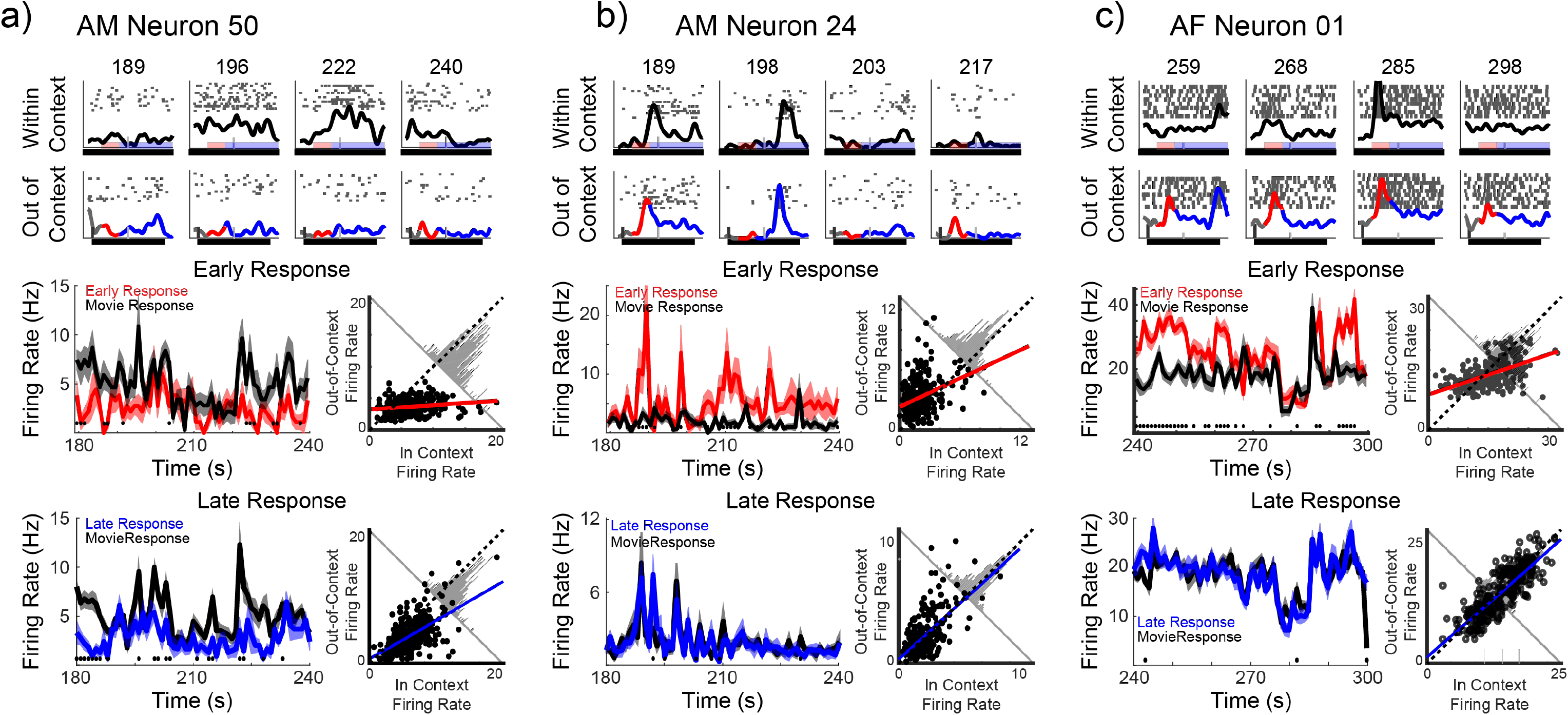
A) Example of a neuron strongly affected by stimulus context. Top row depicts representative spiking responses for four time points to the within context movie data. Each row of spikes represents a movie presentation. The solid black line shows the average spike density function for the given time points. The red bar shows the time frame corresponding to the early response window, and the blue bar shows the late response window. The second row depicts the spiking responses and spike density function (SDF) to the same visual information presented out of context and order. The red region of the SDF shows the early response, and the blue area the late response. The black bar at the bottom shows the stimulus presentation window. The vertical black line shows the stimulus onset. The middle time course shows the reconstructed response to the out-of-context early response (red) and the down sampled in-context response (black) for 60 sec of the full movie. The black dots show time points of significant differences in their firing rate. The scatter plot depicts the correspondence between the in- and out-of-context responses for all 300 snippets. The red line shows the projected linear regression of the data. The histogram shows the density of points when the two contexts firing rates are subtracted from each other. The bottom row displays the late response (blue) in the same format as the early response data. (B) and (C) depict example neurons from the AM and AF face patches, respectively, that were minimally influenced by context, shown in the same format as (A).

**Figure S3.**
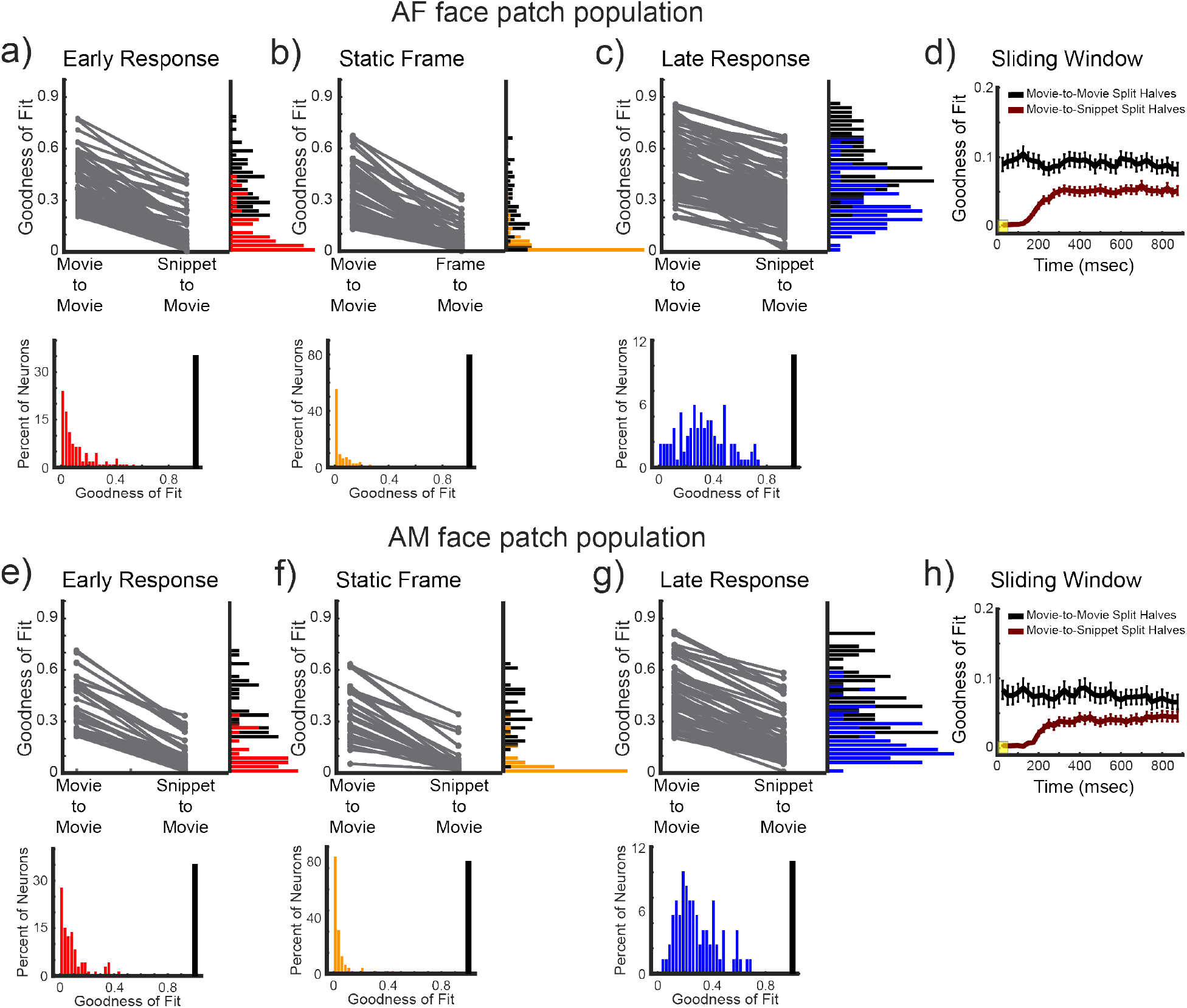
Separate regional population results for the AF and AM face patches. (A) through (D) show the results of the paired comparison of within context and between context goodness-of-fits (r^2^) for the AF population. Whereas, (E) through (H) show the same analyses for the AM face patch populations. (A) and (E) show the early response period, (B) and (F) the static frame responses, (C) and (G) the late response period, and finally (D) and (H) the sliding window analyses.

**Figure S4.**
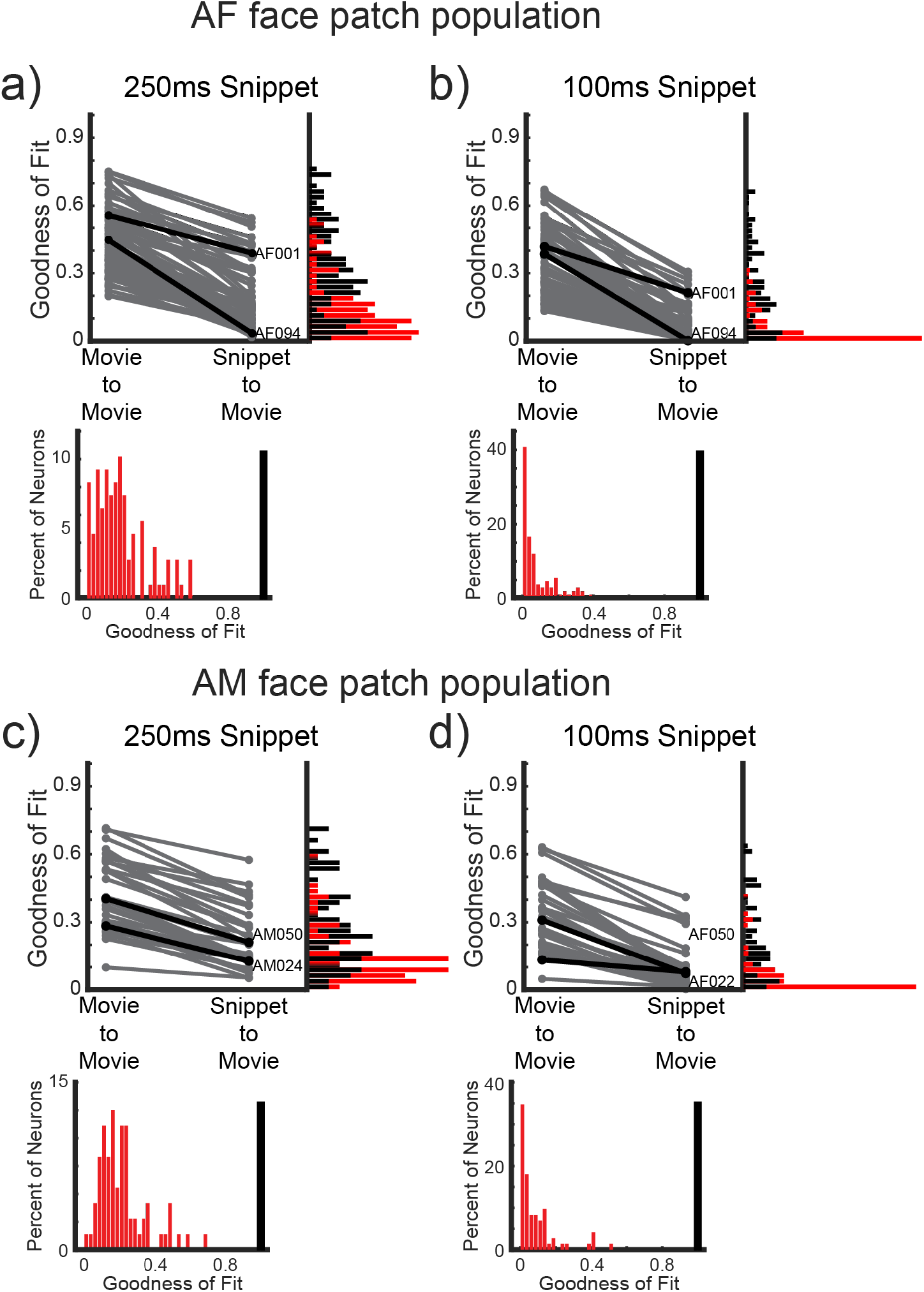
The comparison of the within and between context regressions for the 250 ms snippets and the 100 ms snippets, separated into each face patch population. The data are shown in the same format as **Figures 3**, broken down by region.

**Figure S5.**
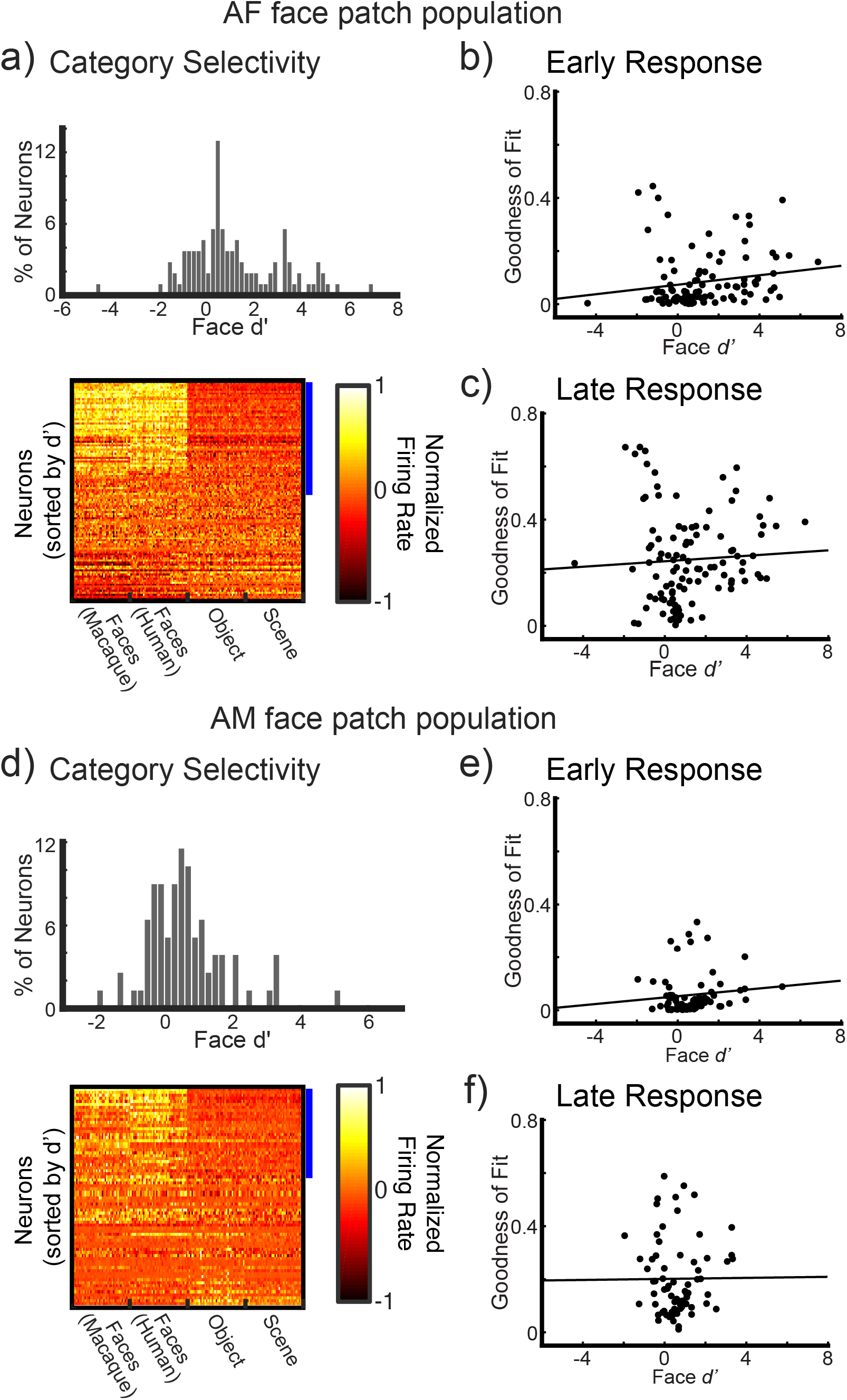
The interaction between face selectivity and context responses, broken down by face patch populations. The data are shown in the same format as **Figure 4**.

## References

Anderson, B., Mruczek, R.E.B., Kawasaki, K., and Sheinberg, D. (2008). Effects of Familiarity on Neural Activity in Monkey Inferior Temporal Lobe. Cereb Cortex 18, 2540–2552.

Aparicio, P.L., Issa, E.B., and DiCarlo, J.J. (2016). Neurophysiological Organization of the Middle Face Patch in Macaque Inferior Temporal Cortex. J Neurosci 36, 12729–12745.

Bartels, A., and Zeki, S. (2004). Functional brain mapping during free viewing of natural scenes. Human Brain Mapping 21, 75–85.

Bell, A.H., Malecek, N.J., Morin, E.L., Hadj-Bouziane, F., Tootell, R.B.H., and Ungerleider, L.G. (2011). Relationship between Functional Magnetic Resonance Imaging-Identified Regions and Neuronal Category Selectivity. J Neurosci 31, 12229–12240.

Brainard, D.H. (1997). The Psychophysics Toolbox. Spatial Vision 10, 433–436.

Chaure, F.J., Rey, H.G., and Quiroga, R.Q. (2018). A novel and fully automatic spike-sorting implementation with variable number of features. Journal of Neurophysiology 120, 1859–1871.

Çukur, T., Nishimoto, S., Huth, A.G., and Gallant, J.L. (2013). Attention during natural vision warps semantic representation across the human brain. Nature Neuroscience 16, 763–770.

Desimone, R., Albright, T., Gross, C., and Bruce, C. (1984). Stimulus-selective properties of inferior temporal neurons in the macaque. J Neurosci 4, 2051–2062.

DiCarlo, J.J., and Maunsell, J.H. (2000). Form representation in monkey inferotemporal cortex is virtually unaltered by free viewing. Nature Neuroscience 3, 814–821.

DiCarlo, J.J., and Maunsell, J.H.R. (2003). Anterior inferotemporal neurons of monkeys engaged in object recognition can be highly sensitive to object retinal position. Journal of Neurophysiology 89, 3264–3278.

Efron, R. (1967). THE DURATION OF THE PRESENT. Ann Ny Acad Sci 138, 713–729.

Franchak, J.M., Heeger, D.J., Hasson, U., and Adolph, K.E. (2016). Free Viewing Gaze Behavior in Infants and Adults. Infancy 21, 262–287.

Freiwald, W.A., Tsao, D.Y., and Livingstone, M.S. (2009). A face feature space in the macaque temporal lobe. Nature Neuroscience 12, 1187–1196.

Furl, N., Hadj-Bouziane, F., Liu, N., Averbeck, B.B., and Ungerleider, L.G. (2012). Dynamic and Static Facial Expressions Decoded from Motion-Sensitive Areas in the Macaque Monkey. J Neurosci 32, 15952–15962.

Gross, C.G., Rocha-Miranda, C.E., and Bender, D.B. (1972). Visual properties of neurons in inferotemporal cortex of the Macaque. Journal of Neurophysiology 35, 96–111.

Hasson, U., and Honey, C.J. (2012). Future trends in Neuroimaging: Neural processes as expressed within real-life contexts. NeuroImage 62, 1272–1278.

Hasson, U., Nir, Y., Levy, I., Fuhrmann, G., and Malach, R. (2004). Intersubject Synchronization of Cortical Activity During Natural Vision. Science 303, 1634–1640.

Hasson, U., Yang, E., Vallines, I., Heeger, D.J., and Rubin, N. (2008). A Hierarchy of Temporal Receptive Windows in Human Cortex. J Neurosci 28, 2539–2550.

Hasson, U., Malach, R., and Heeger, D.J. (2010). Reliability of cortical activity during natural stimulation. Trends in Cognitive Sciences 14, 40–48.

Henderson, J.M. (2003). Human gaze control during real-world scene perception. Trends Cogn Sci 7, 498–504.

Hung, C.P., Kreiman, G., Poggio, T., and DiCarlo, J.J. (2005). Fast Readout of Object Identity from Macaque Inferior Temporal Cortex. Science 310, 863–866.

Huth, A.G., Nishimoto, S., Vu, A.T., and Gallant, J.L. (2012). A continuous semantic space describes the representation of thousands of object and action categories across the human brain. Neuron 76, 1210–1224.

Jellema, T., and Perrett, D.I. (2003). Perceptual History Influences Neural Responses to Face and Body Postures. J Cognitive Neurosci 15, 961–971.

Jellema, T., Maassen, G., and Perrett, D.I. (2004). Single Cell Integration of Animate Form, Motion and Location in the Superior Temporal Cortex of the Macaque Monkey. Cereb Cortex 14, 781–790.

Keysers, C., and Perrett, D.I. (2004). Demystifying social cognition: a Hebbian perspective. Trends Cogn Sci 8, 501–507.

Kourtzi, Z., and DiCarlo, J.J. (2006). Learning and neural plasticity in visual object recognition. Curr Opin Neurobiol 16, 152–158.

Kuravi, P., and Vogels, R. (2017). Effect of adapter duration on repetition suppression in inferior temporal cortex. Sci Rep-Uk 7, 3162.

Leopold, D.A., and Park, S.H. (2020). Studying the visual brain in its natural rhythm. NeuroImage 116790.

Leszczynski, M., and Schroeder, C.E. (2019). The Role of Neuronal Oscillations in Visual Active Sensing. Frontiers in Integrative Neuroscience 13, 32.

Mantini, D., Hasson, U., Betti, V., Perrucci, M.G., Romani, G.L., Corbetta, M., Orban, G.A., and Vanduffel, W. (2012). Interspecies activity correlations reveal functional correspondence between monkey and human brain areas. Nat Methods 9, 277–282.

Mantini, D., Corbetta, M., Romani, G.L., Orban, G.A., and Vanduffel, W. (2013). Evolutionarily Novel Functional Networks in the Human Brain? J Neurosci 33, 3259–3275.

McMahon, D.B.T., Bondar, I.V., Afuwape, O.A.T., Ide, D.C., and Leopold, D.A. (2014a). One month in the life of a neuron: longitudinal single-unit electrophysiology in the monkey visual system. Journal of Neurophysiology 112, 1748–1762.

McMahon, D.B.T., Jones, A.P., Bondar, I.V., and Leopold, D.A. (2014b). Face-selective neurons maintain consistent visual responses across months. Proceedings of the National Academy of Sciences 111, 8251–8256.

McMahon, D.B.T., Russ, B.E., Elnaiem, H.D., Kurnikova, A.I., and Leopold, D.A. (2015). Single-Unit Activity during Natural Vision: Diversity, Consistency, and Spatial Sensitivity among AF Face Patch Neurons. J Neurosci 35, 5537–5548.

Melcher, D. (2001). Persistence of visual memory for scenes. Nature 412, 401–401.

Melcher, D., and Colby, C.L. (2008). Trans-saccadic perception. Trends Cogn Sci 12, 466–473.

Meyer, T., and Olson, C.R. (2011). Statistical learning of visual transitions in monkey inferotemporal cortex. Proc National Acad Sci 108, 19401–19406.

Meyer, T., Ramachandran, S., and Olson, C.R. (2014a). Statistical Learning of Serial Visual Transitions by Neurons in Monkey Inferotemporal Cortex. J Neurosci 34, 9332–9337.

Meyer, T., Walker, C., Cho, R.Y., and Olson, C.R. (2014b). Image familiarization sharpens response dynamics of neurons in inferotemporal cortex. Nat Neurosci 17, 1388–1394.

Miller, E., Li, L., and Desimone, R. (1991). A neural mechanism for working and recognition memory in inferior temporal cortex. Science 254, 1377–1379.

Mitchell, J.F., and Leopold, D.A. (2015). The marmoset monkey as a model for visual neuroscience. Neurosci Res 93, 20–46.

Miyashita, Y. (1988). Neuronal correlate of visual associative long-term memory in the primate temporal cortex. Nature 335, 817–820.

Mosher, C.P., Zimmerman, P.E., and Gothard, K.M. (2014). Neurons in the Monkey Amygdala Detect Eye Contact during Naturalistic Social Interactions. Curr Biol 24, 2459–2464.

Müller, J.R., Metha, A.B., Krauskopf, J., and Lennie, P. (2001). Information Conveyed by Onset Transients in Responses of Striate Cortical Neurons. J Neurosci 21, 6978–6990.

Nelissen, K., Luppino, G., Vanduffel, W., Rizzolatti, G., and Orban, G.A. (2005). Observing Others: Multiple Action Representation in the Frontal Lobe. Science 310, 332–336.

Nelissen, K., Vanduffel, W., and Orban, G.A. (2006). Charting the Lower Superior Temporal Region, a New Motion-Sensitive Region in Monkey Superior Temporal Sulcus. J Neurosci 26, 5929–5947.

Nelissen, K., Borra, E., Gerbella, M., Rozzi, S., Luppino, G., Vanduffel, W., Rizzolatti, G., and Orban, G.A. (2011). Action Observation Circuits in the Macaque Monkey Cortex. J Neurosci 31, 3743–3756.

Oram, M.W., and Perrett, D.I. (1992). Time course of neural responses discriminating different views of the face and head. J Neurophysiol 68, 70–84.

Park, S.H., Russ, B.E., McMahon, D.B.T., Koyano, K.W., Berman, R.A., and Leopold, D.A. (2017). Functional Subpopulations of Neurons in a Macaque Face Patch Revealed by Single-Unit fMRI Mapping. Neuron 95, 971–981.e5.

Park, S.H., Koyano, K.W., Russ, B.E., Waidmann, E.N., McMahon, D.B.T., and Leopold, D.A. (2021). Parallel functional subnetworks embedded in the macaque face patch system. Biorxiv 2021.10.25.465775.

Perrett, D.I., Hietanen, J.K., Oram, M.W., and Benson, P.J. (1992). Organization and functions of cells responsive to faces in the temporal cortex. Philosophical Transactions of the Royal Society of London Series B, Biological Sciences 335, 23–30.

Perrett, D.I., Xiao, D., Barraclough, N.E., Keysers, C., and Oram, M.W. (2009). Seeing the future: Natural image sequences produce “anticipatory” neuronal activity and bias perceptual report. Q J Exp Psychol 62, 2081–2104.

Polosecki, P., Moeller, S., Schweers, N., Romanski, L.M., Tsao, D.Y., and Freiwald, W.A. (2013). Faces in Motion: Selectivity of Macaque and Human Face Processing Areas for Dynamic Stimuli. J Neurosci 33, 11768–11773.

Popivanov, I.D., Jastorff, J., Vanduffel, W., and Vogels, R. (2014). Heterogeneous Single-Unit Selectivity in an fMRI-Defined Body-Selective Patch. J Neurosci 34, 95–111.

Quiroga, R.Q., Nadasdy, Z., and Ben-Shaul, Y. (2004). Unsupervised spike detection and sorting with wavelets and superparamagnetic clustering. Neural Computation 16, 1661–1687.

Ramachandran, S., Meyer, T., and Olson, C.R. (2017). Prediction suppression and surprise enhancement in monkey inferotemporal cortex. J Neurophysiol 118, 374–382.

Rolls, E.T., Baylis, G.C., Hasselmo, M.E., and Nalwa, V. (1989). The effect of learning on the face selective responses of neurons in the cortex in the superior temporal sulcus of the monkey. Exp Brain Res 76, 153–164.

Russ, B.E., and Leopold, D.A. (2015). Functional MRI mapping of dynamic visual features during natural viewing in the macaque. NeuroImage 109, 84–94.

Russ, B.E., Kaneko, T., Saleem, K.S., Berman, R.A., and Leopold, D.A. (2016). Distinct fMRI Responses to Self-Induced versus Stimulus Motion during Free Viewing in the Macaque. J Neurosci 36, 9580–9589.

Schiller, P.H., and Malpeli, J.G. (1977). Properties and tectal projections of monkey retinal ganglion cells. J Neurophysiol 40, 1443–1443.

Schroeder, C.E., Wilson, D.A., Radman, T., Scharfman, H., and Lakatos, P. (2010). Dynamics of Active Sensing and perceptual selection. Curr Opin Neurobiol 20, 172–176.

Schwiedrzik, C.M., and Freiwald, W.A. (2017). High-Level Prediction Signals in a Low-Level Area of the Macaque Face-Processing Hierarchy. Neuron 96, 89–97.e4.

Sheinberg, D.L., and Logothetis, N.K. (2001). Noticing Familiar Objects in Real World Scenes: The Role of Temporal Cortical Neurons in Natural Vision. J Neurosci 21, 1340–1350.

Shepherd, S.V., Steckenfinger, S.A., Hasson, U., and Ghazanfar, A.A. (2010). Human-Monkey Gaze Correlations Reveal Convergent and Divergent Patterns of Movie Viewing. Curr Biol 20, 649–656.

Singer, J.M., and Sheinberg, D.L. (2010). Temporal Cortex Neurons Encode Articulated Actions as Slow Sequences of Integrated Poses. J Neurosci 30, 3133–3145.

Sliwa, J., and Freiwald, W.A. (2017). A dedicated network for social interaction processing in the primate brain. Science 356, 745–749.

Tsao, D.Y., Freiwald, W.A., Tootell, R.B.H., and Livingstone, M.S. (2006). A Cortical Region Consisting Entirely of Face-Selective Cells. Science 311, 670–674.

Vanderwal, T., Kelly, C., Eilbott, J., Mayes, L.C., and Castellanos, F.X. (2015). Inscapes: A movie paradigm to improve compliance in functional magnetic resonance imaging. Neuroimage 122, 222–232.

Vangeneugden, J., Pollick, F., and Vogels, R. (2009). Functional Differentiation of Macaque Visual Temporal Cortical Neurons Using a Parametric Action Space. Cereb Cortex 19, 593–611.

Vinken, K., Beeck, H.O. de, and Vogels, R. (2018). Face Repetition Probability Does Not Affect Repetition Suppression in Macaque Inferotemporal Cortex. J Neurosci 38, 0462–18.

Vodrahalli, K., Chen, P.-H., Liang, Y., Baldassano, C., Chen, J., Yong, E., Honey, C., Hasson, U., Ramadge, P., Norman, K.A., et al. (2017). Mapping between fMRI responses to movies and their natural language annotations. NeuroImage 1–9.

Wang, H.X., Freeman, J., Merriam, E.P., Hasson, U., and Heeger, D.J. (2012). Temporal eye movement strategies during naturalistic viewing. Journal of Vision 12, 16.

Woloszyn, L., and Sheinberg, D.L. (2012). Effects of long-term visual experience on responses of distinct classes of single units in inferior temporal cortex. Neuron 74, 193–205.

Wurtz, R.H., Joiner, W.M., and Berman, R.A. (2011). Neuronal mechanisms for visual stability: progress and problems. Philosophical Transactions of the Royal Society B: Biological Sciences 366, 492–503.

